# Refining liquid chromatography conditions across platforms to democratize single-cell proteomics

**DOI:** 10.64898/2025.12.16.694739

**Authors:** Yiwen Wang, Qiuye Zhao, William K.M. Lai, Haiyuan Yu

## Abstract

Single-cell proteomics has emerged as a powerful approach for characterizing cellular heterogeneity. Here, we present an optimized scProteomic workflow that enhances proteome coverage and quantification by refining liquid chromatography (LC) conditions across platforms (both Evosep and nanoElute2). Importantly, our results show that using our optimized LC conditions, even with Bruker timsTOF HT, a machine not designed for scProteomics applications, we achieved solid performance with single cell samples that allows meaningful biological discoveries. First, we compared power to detect differentially-expressed genes/proteins between scRNA-seq and scProteomics at single cell level, and demonstrated that scProteomics showed an excellent performance at small sample sizes. To further validate this finding, we applied our optimized workflow to study the proteomic response to oxidative stress using a limited number of single cells. Our scProteomics results successfully detected 3 distinct cell populations, reflecting correctly the 3 cell lines used, and captured the dysregulation of antioxidant enzymes, molecular chaperones and ubiquitin-proteasome system, reflecting a multi-faceted response to oxidative stress that is uniform across distinct cell lines. Our results highlighted the potential of scProteomics to resolve subtle perturbations and provide a readout of cellular states for the broader community, including users operating non-scProteomics-dedicated machines.

## Introduction

Liquid chromatography and mass spectrometry–based single-cell proteomics (scProteomics) has rapidly evolved in recent years, achieving significant progress in proteome identification and quantification through efforts to optimize sample preparation^1,2^, LC column and gradient^3^, MS data acquisition^4^ and spectral library^5,6^.

Beyond technical advances, several studies have compared scProteomic data with matched or reference scRNA-seq datasets to examine the cross-modality correlation^7,8^, sampling efficiency^9^, covariation^10^, variance and noise level^11^. These studies have revealed that scProteomics allows for an accurate measure of cell state, complementing scRNA-seq by capturing regulatory events beyond mRNA abundance. This underscores the unique advantage of scProteomics in resolving direct functional cellular behaviors.

While most studies have demonstrated that scProteomics can robustly distinguish cell types^9^ and cell states^12^ typically defined by the differences across hundreds of proteins, it remains challenging to resolve short-term perturbations involving only a limited subset of dysregulated proteins. For example, upon oxidative stress, cells activate multiple defenses to restore redox homeostasis. These involve enzymatic antioxidants like superoxide dismutases (SODs), catalases (CAT), glutathione peroxidases (GPXs) and peroxiredoxins (PRDXs) for detoxification, redox-sensitive transcription factors to induce stress responsive gene, molecular chaperones to refold misfolded proteins, and proteasomes to degrade damaged proteins^13^.

scProteomics workflows often require several dedicated equipment at different steps, such as dedicated single cell sorter^14^, automated nanoliter pipettor^15^ and various microfluidic devices^12,16^ for sample preparation. More recently, scProteomics-dedicated mass spectrometer machines have been developed, such as Bruker timsTOF Ultra2 and Thermo Orbitrap Astral. However, most biology laboratories do not have access to many of the dedicated equipment, limiting the access to the scProteomic technology. In addition to instrumentation, the inherently low throughput of scProteomics restricts its biological applicability. Unlike scRNA-seq, which routinely profiles thousands to tens of thousands of cells per run, scProteomics throughput remains far more limited—even with the development of various multiplexing strategies^17,18^. These multiplexing methods often rely on chemical labeling reagents, increase experimental cost, and can increase acquisition stochasticity.

In this study, we evaluated different LC platforms (namely, Evosep and nanoElute2 with fundamentally different design principles) and optimized LC conditions for scProteomics. We aim to develop a workflow without requiring dedicated equipment or large sample sizes, which allows many more laboratories to take advantage of the scProteomics technology for their research projects. All of our single cell samples were processed using a non-scProteomics-dedicated timsTOF HT machine. And, we applied our optimized workflow to dissect the single-cell oxidative stress response at small sample sizes. Despite only profiling a limited number of single cells per condition, we demonstrated that scProteomics can sensitively capture proteomic remodeling underlying redox adaptation and resolve subpopulations corresponding to the underlying cell identities. These results highlight that, upon oxidative stress, intrinsic cell identity dominates proteomic variation more than the uniform stress-induced changes, underscoring the capacity of scProteomics to reveal subtle and biologically relevant changes even with limited sample sizes.

## Results

### Enhanced Proteome Coverage and Quantification Through Optimized LC conditions

To optimize LC conditions for single-cell level characterization, we compared between different LC systems, and across different LC columns on timsTOF-HT. Specifically, we tested two LC platforms: nanoElute2, which offered high flexibility in gradient and flow rate customization, and Evosep, which provided pre-defined LC methods optimized for throughput and reproducibility. While Evosep enabled higher sample throughput and reduced carryover, nanoElute2 offered complete customization of flow rate and gradient. We tested both commercially available columns commonly used in scProteomics and our own homemade columns, which allow greater flexibility in key parameters such as inner diameter (75μm and 50μm), column length (15cm, 10cm and 5cm), and C18 particle size (1.9μm, 1.7μm and 1.5μm).

On Evosep, columns with smaller particle size gave better proteome coverage and quantification at 3 ng of peptide input (**Figure 1**a, b). While smaller particle size enhanced separation and therefore proteome identification and quantification, it also increased backpressure, limiting column length and throughput. Our results show that both PepSep and IonOpticks commercial columns performed very similarly. Our homemade column, with a larger particle size (1.9μm), detected slightly less, but still solid numbers of proteins with both low-input (**Figure 1** c, d) and single-cell (**Figure 1** c, d) samples.

**Figure 1.**
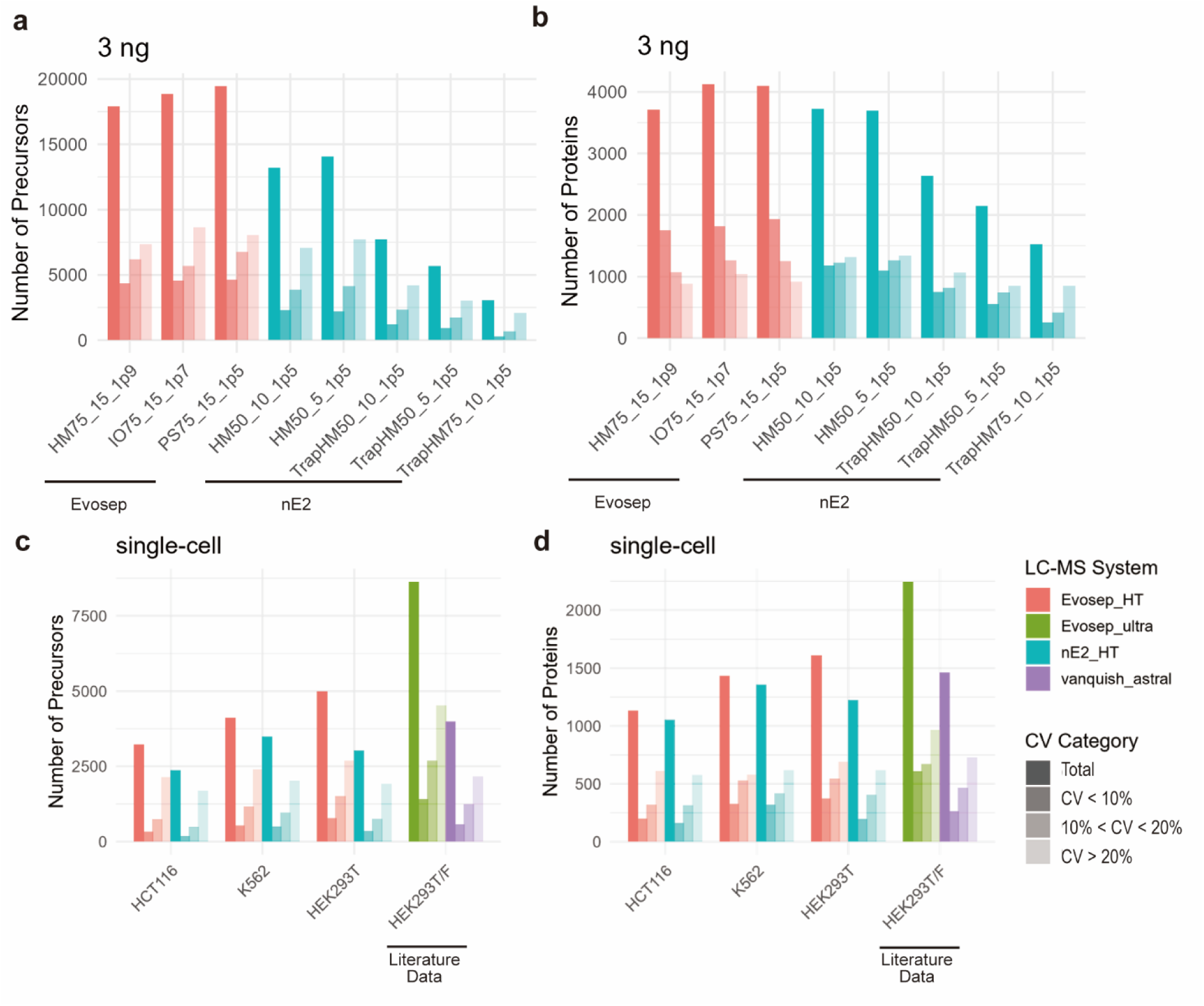
Optimization of scProteomics workflow. a) Precursor identification at 3ng of peptide input on different columns and different LC systems. HM50_10_1p5: Homemade column 50μm ID, 10cm length,1.5μm C18; HM50_5_1p5: Homemade column 50μm ID, 5cm length, 1.5μm C18; HM75_10_1p5: Homemade column 75μm ID, 10cm length, 1.5μm C1; HM75_15_1p9: Homemade column 75μm ID, 15cm length, 1.9μm C18; IO75_15_1p7: IonOpticks column 75μm ID, 15cm length, 1.7μm C18; PS75_15_1p5: PepSep column 75μm ID, 15cm length, 1.5μm C1. b) Protein identification at 3ng of peptide input on different columns and different LC systems. c) Precursor identification at single-cell input on different LC-MS systems. d) Protein identification at single-cell input on different LC-MS systems. The first bar represents the number of precursors found in three technical replicates, the second bar those with CV<10% and the third bar those with 10%<CV<20%.

On nanoElute2, one-column separation outperformed two-column separation with more than 40% increase in protein identification (**Figure 1**a, b). Although two-column configuration accelerated sample loading and improved throughput, it also introduced additional dead volume and potential sample loss, leading to broader peaks and reduced sensitivity. Therefore, the one-column setup was especially advantageous for low-input, especially single-cell samples using nanoElute2. We further tested several column setups with different combinations of lengths, inner diameters, and particle sizes. Our results show that the 50μm column at 5cm length filled with 1.5μm C18 particles (HM50-5-1p5) has the highest identification at both precursor and protein levels.

Using the Evosep coupled with our non-scProteomics-dedicated timsTOF HT machine, we quantified from a single HEK293T cell 5,000 precursors and 1,600 proteins, among which more than 50% of proteins showed CV less than 20 (**Figure 1** c, d). We compared our scProteomics results on timsTOF HT with the published data using state-of-the-art scProteomics-dedicated MS instruments and similar cell lines^12,19^. While we observed an approximate 25% increase in protein identification using Evosep coupled with timsTOF Ultra than timsTOF HT, we note that the performance using timsTOF HT is comparable with that of using Thermo Orbitrap Astral. Therefore, we believe we have established an optimized scProteomics workflow that can be implemented without dedicated equipment, and even with a non-scProteomics-dedicated timsTOF HT can detect >1000 proteins robustly across diverse cell types. Our workflow can be integrated into many research projects and help lead to meaningful biological discoveries.

### Comparison Between Single-cell Proteome and Transcriptome

Next, we compared our K562 single-cell proteome produced by Evosep and timsTOF HT with a reference K562 Drop-seq dataset^20^. Analysis of the 2,146 shared genes between the two modalities showed that scProteomics exhibited much higher pairwise correlation than scRNAseq (**Figure 2**a-c). This was likely because proteins exist at much higher copy numbers than their corresponding transcripts in each cell^21^, making them less affected by Poisson sampling noise, PCR amplification bias and reflecting stronger biological covariation across cells. This also demonstrated that even with a non-scProteomics-dedicated machine, our measurements at single cell level are highly reproducible from cell to cell.

**Figure 2.**
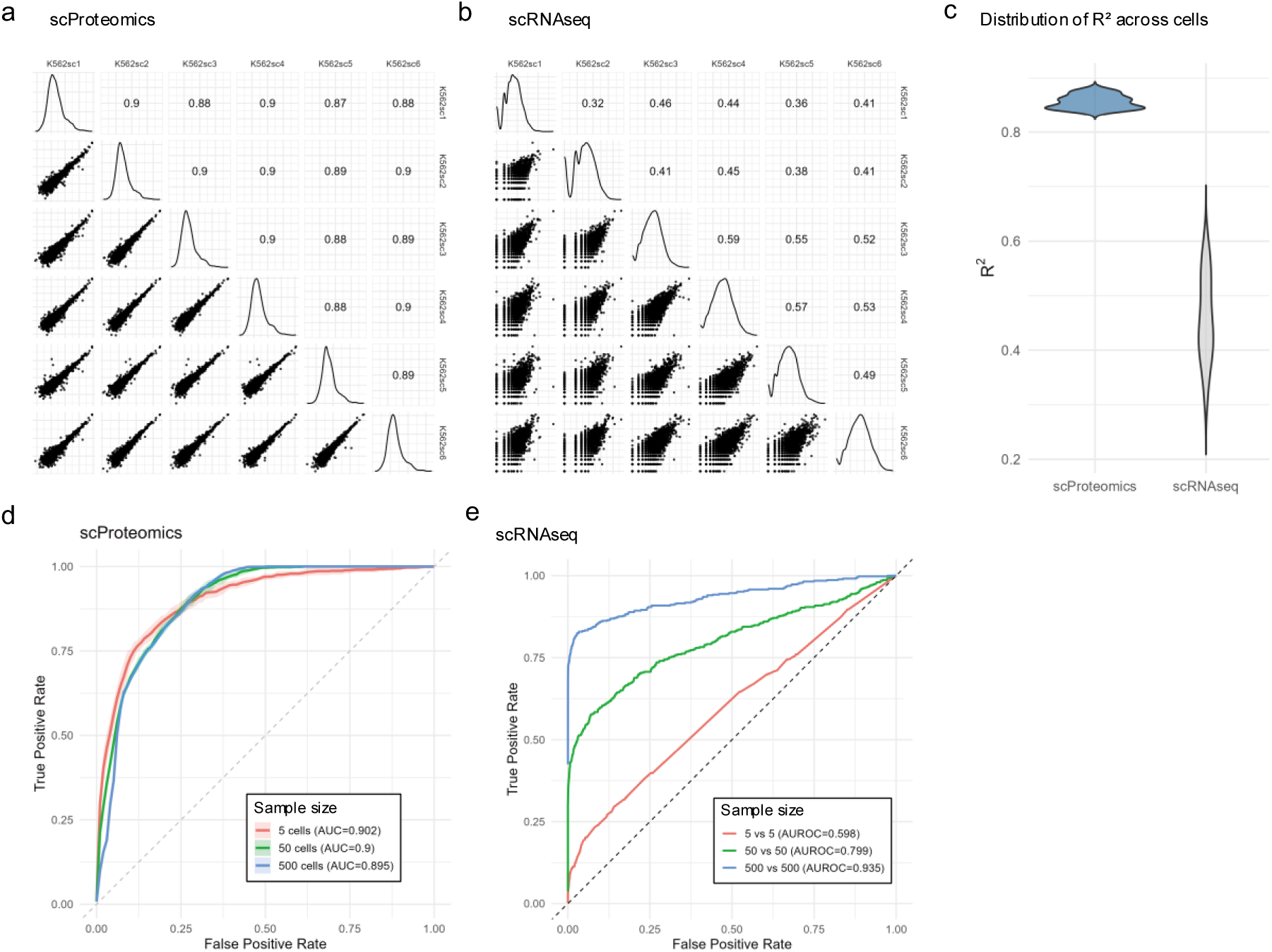
Comparative analysis of scProteomics and scRNAseq. a) Scatter plot of cell–cell correlations across individual cells measured by scProteomics. b) Scatter plot of cell–cell correlations across individual cells measured by scRNA-seq. c) R^2^ distribution across cells. d) ROC curve of scProteomics at different sample sizes. e) ROC curve of scRNA-seq at different sample sizes.

We reasoned that the lower variance in scProteomics would significantly enhance our ability to detect differentially expressed (DE) proteins with much fewer number of cells. This is important, because the throughput of scProteomics is inherently much more limited and significantly lower throughput than scRNA-seq. To evaluate this, we conducted DE power analysis through simulations using gene/protein-specific expression means, variances, and detection probabilities that are measured in the experimental scProteomics and scRNA-seq datasets (Methods). In these simulations, true DE genes or proteins may be missed either because of high variance or because of low expression probability. Our results show that scRNA-seq is more susceptible to high variability at small sample sizes and only begins to exhibit robust DE power when profiling reached ∼1,000 single cells, at which point gene-level mean and variance estimates stabilized and sampling noise was effectively averaged out^22^ (**Figure 2**e). By contrast, scProteomics achieves its best performance at smallest sample sizes (**Figure 2**d), effectively capturing the most readily detectable DE signals. As sample size increases, more proteins with low expression probability enter the analysis: while this allows more true positives to be recovered, the high sparsity also introduces additional errors, compromising the overall performance. For scProteomics, sensitivity rather than sample size is the key driver of DE power.

### scProteomics Dissects Oxidative Stress Response

To assess the ability of our scProteomics workflow to detect biological stress responses, we examined the oxidative stress response in three cell lines (K562, HEK293T, and hct116) under acute hydrogen peroxide treatment (300 uM) for 30 minutes^23^ and detected changes in single-cell proteomes using Evosep and timsTOF HT. Guided by this DE power analysis, and given the homogeneity of each cell line, we profiled about five single cells per line and treatment to ensure sufficient DE power while minimizing unnecessary sampling First, we performed UMAP analysis on the stress response dataset and observed strong separation across different cell lines and weaker separation upon treatment (**Figure 3**a). This pattern reflected the dominance of cell-type-specific expression programs, while the short-term drug treatment induced relatively subtle proteomic changes that did not override the intrinsic differences between cell types.

**Figure 3.**
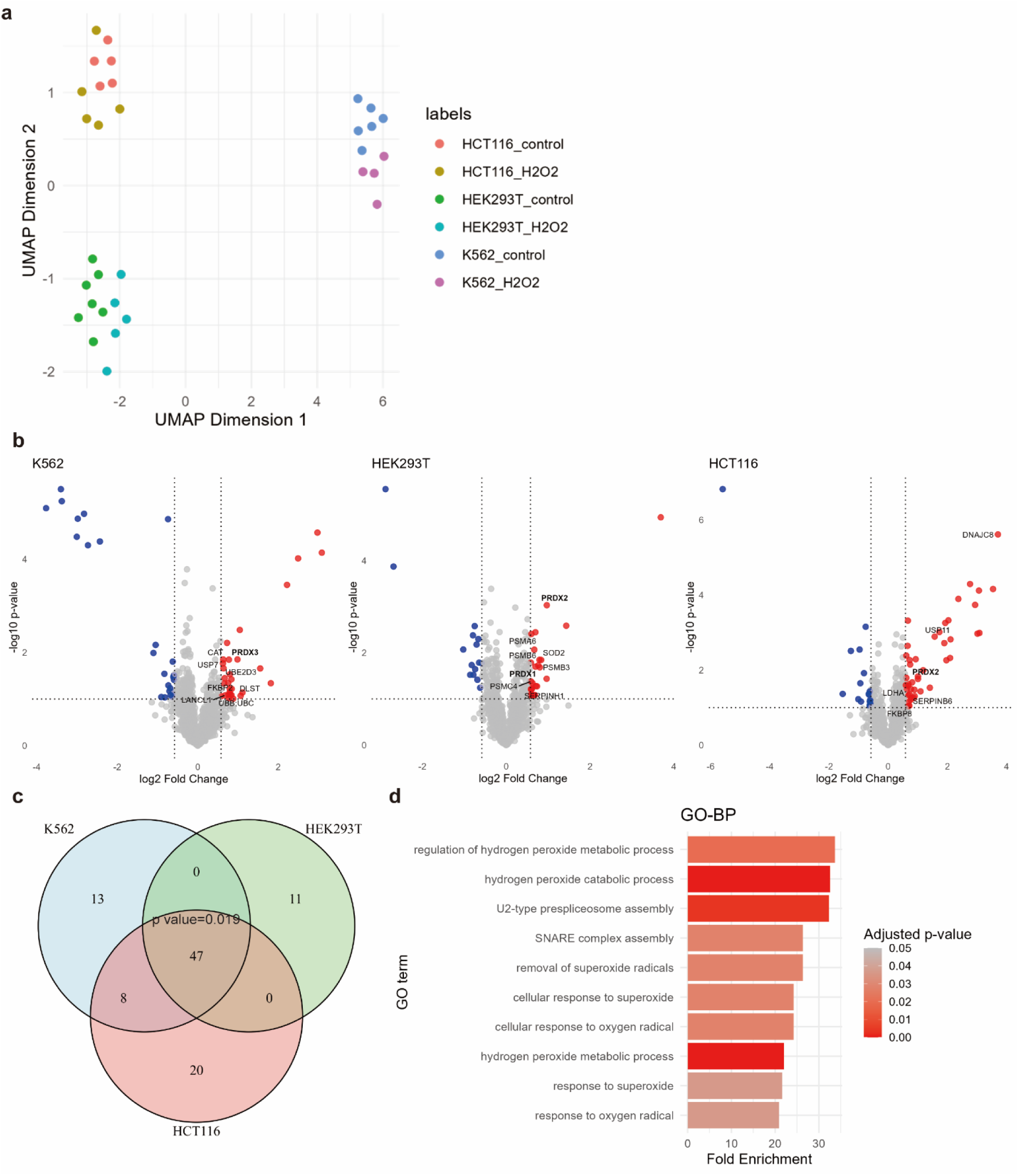
scProteomics captures oxidative stress response. a) UMAP plot of single cells upon H_2_O_2_ treatment. b) Volcano plot of quantitative protein differences between treated and untreated single cells. c) Venn diagram showing the overlap of the up-regulated proteins within the interactome network. An up-regulated protein in one cell line is considered to be detected in another cell line if itself or its direct interactor in the interactome is up-regulated in the other cell line upon oxidative stress. d) Bar plot of top ten enriched GO BP terms among up-regulated proteins.

We next investigated the DE proteins between the treatment and control group within each cell line. Among the significantly upregulated proteins, there were a large proportion of known oxidative stress responders (**Figure 3**b). These included canonical antioxidant such as SOD, PRDXs, and CAT, which constituted the primary enzymatic defense against reactive oxygen species. In addition, molecular chaperones (e.g., DNAJ, FKBP, SERPIN) and components in proteasome and ubiquitin system were upregulated, reflecting protein quality control activity^13^. Among them, proteins with high basal abundance such as PRDX family members and components in proteasome and ubiquitin system were consistently differentially expressed across all three cell lines, suggesting a shared oxidative stress–responsive mechanism that underlies potential cell-type-specific adaptations and underscoring their essential roles in maintaining redox balance and proteome integrity under stress conditions. Across the three cell lines, proteins up-regulated upon oxidative stress showed a significant overlap across the 3 cell lines (**Figure 3**c). Together, these changes highlighted a coordinated cellular program that coupled ROS detoxification with proteome stabilization upon oxidative stress (**Figure 3** d). These observations confirmed that scProteomics, using our optimized workflow without dedicated scProteomic equipment and processed by non-scProteomics-dedicated timsTOF HT, could recapitulate many stress responsive proteins and pathways, despite reduced depth and quantification compared to bulk proteome measurements.

## Discussion

By evaluating column configurations on two different LC platforms, we found that each system requires a different optimal configuration. Our results highlighted the necessity of optimization tailored to specific LC systems. The nanoElute2 system favored a single-column configuration without using a trap column and with low inner diameter (50μm) and low flow rate (100 nL/min), which minimized dead volume, improved peak sharpness, minimized in-column adsorption loss, and maximized ionization efficiency. In contrast, the Evosep system performed best with its standardized trap-and-elute workflow with small particle size (1.5μm), which maximized reproducibility, and improved peak capacity and signal-to-noise ratio. These differences reflected the fundamentally distinct design philosophies of the two systems: nanoElute2 provided flexibility and ultra-low-flow control ideal for customized applications, whereas Evosep offered robustness and consistency optimized for large-scale, high-throughput studies.

Through DE power analysis, we found that scProteomics did not necessarily benefit from large sample sizes in the case of a homogeneous population. The results highlighted the difference between single-cell proteomics and transcriptomics measurements. While scRNA-seq benefited substantially from large sample sizes that stabilize variance estimates and rescue low-abundance transcripts, scProteomics derived its statistical power primarily from detection sensitivity rather than sampling depth. The higher measurement precision in scProteomics allowed for reliable identification of high-abundance DE proteins even with limited cell numbers. This highlighted that, for scProteomics, improvement in ion sampling would be more critical than increasing cell throughput for enhancing biological resolution. Importantly, this demonstrated that scProteomics can be effectively powered even with relatively small sample sizes, lowering the barrier to entry and helping democratize scProteomic studies for laboratories without access to ultra-high-throughput platforms.

By applying our optimized scProteomic workflow to an acute oxidative stress experiment, we showcased that scProteomics could resolve stress-induced dysregulation beyond cell-type-specific expression. Finally, we emphasize that while our scProteomics workflow clearly detected the stress response within single cells, it also enabled us to distinctly recognize three subpopulations of cells (corresponding to the 3 different cell lines used) within our treated samples. Therefore, our optimized scProteomics workflow, even with non-scProteomics-dedicated mass spectrometers, can enable users to resolve biologically meaningful subpopulations and can help address key questions from a wide range of biological and clinical samples.

## Conclusion

Through systematic optimization of LC configurations, we established a robust single-cell proteomics workflow without special equipment and across LC platforms. We demonstrated that LC performance was highly system-dependent and required tailored optimization to balance sensitivity, throughput, and reproducibility. Comparative analysis between scProteomics and scRNA-seq further highlighted the distinct statistical and biological properties of the two modalities: scProteomics exhibited higher pairwise correlation and lower variance, allowing for a significantly improved DE power even at small sample sizes. Finally, application of this workflow to oxidative stress perturbation revealed that scProteomics, even with non-scProteomics-dedicated equipment, could capture coordinated cellular responses, such as redox detoxification and proteome stabilization, while detecting three distinct cell populations in the meantime, despite limited proteome coverage. Together, these findings demonstrated the potential of optimized single-cell proteomics to bridge quantitative precision with biological resolution, enabling deeper insights into cellular states and dynamic regulatory mechanisms at the proteome level.

### Experimental Section

#### Cell Culture and Treatment

K562 cells were cultured in Iscove’s Modified Dulbecco’s Medium (IMDM). HCT116 cells were cultured in Dulbecco’s Modified Eagle Medium/Nutrient Mixture F-12 (DMEM/F-12). HEK293T cells were cultured in Dulbecco’s Modified Eagle Medium (DMEM). The cells were incubated in serum-free and pyruvate-free DMEM medium containing 0.3 mM H2O2 (Sigma-Aldrich, H1009) for 30 min at 37℃. The cells were harvested and then subjected to ice-cold PBS washes before DAPI-negative single live cell FACS sorting on Sony MA900 sorter using single-cell (3-drop) mode.

#### Single-cell Sample Preparation

Cells were sorted into lobind tubes (Eppendorf, 22431081) containing 2 uL of miliQ water, briefly spun down, frozen on dry ice and stored at -80℃. Cells were lysed by 3 cycles of freeze in dry ice ethanol bath and thaw at 37℃ followed by the addition of 1 uL of 0.1% DDM in 50 mM Triethylammonium bicarbonate (TEAB) and 3 min sonication. The lysate was digested by the addition of 1 uL of 2 ng lysC-trypsin mix and incubation at 37℃ for 4h. The digest was acidified by the addition of 5 uL of 1% FA and centrifuged at 21130 g, 4℃ for 15 min. The supernatant was either kept in silanized glass vials for direct injection to nanoElute2 or loaded on evotips.

#### LC-MS/MS analysis

Proteomics analysis was conducted using diaPASEF on timsTOF-HT coupled with nanoElute2 or Evosep One. Diluted peptide samples were separated using different C18 columns: Homemade column HM50_10_1p5 (50μm ID, 10cm length,1.5μm C18), Homemade column HM50_5_1p5 (50μm ID, 5cm length, 1.5μm C18) and Homemade column HM75_10_1p5 (75μm ID, 10cm length, 1.5μm C18) on nanoElute2 using a 30-min gradient and Homemade column HM75_15_1p9 (75μm ID, 15cm length, 1.9μm C18), IonOpticks column IO75_15_1p7 (75μm ID, 15cm length, 1.7μm C18) and PepSep column PS75_15_1p5 (75μm ID, 15cm length, 1.5μm C18) on EvoSep One using the Whisper Zoom 40SPD method. FACS-scProteomics samples were separated using the Homemade column HM50_5_1p5 (50μm ID, 5cm length, 1.5μm C18) on nanoElute2 or using the PepSep column PS75_15_1p5 (75μm ID, 15cm length, 1.5μm C18) on EvoSep One. For the MS parameter, the mobility range was set to 1/K0 0.7 to 1.3 V·s/cm^-2^ and the mass range was set to m/z 300 to 975. The accumulation time was set to 100 ms and high sensitivity mode was enabled.

#### Data Analysis

The single-cell runs were searched against a 10-cell spectral library using DIANN v1.9.2.

For single-cell protein and RNA comparison, 10k Human K562-r Cells, Singleplex Sample (Next GEM) dataset and the K562 single-cell proteome subset were used. For correlation analysis, log2-transformed gene or protein quantification events of two cells were plotted against each other without imputation. For differential expression power analysis, powsimR (v1.2.5) limma-voom mode was used for scRNAseq. Single-cell proteome matrices were simulated by downsampling the expression probability, mean, variance of genes and tested by limma (v3.62.2). A vector of ground-truth log2 fold-changes across 2,000 genes was generated, consisting of 5% DE genes with ∣ log2FC ∣> 1, sampled from a gamma distribution and randomly assigned positive or negative signs, and 95% non-DE genes with ∣ log2FC ∣≤ 1. For each gene, expression values were then simulated separately for two groups (control and treatment). The baseline mean expression and variance for each gene were taken from our experimental scProteomics data, and group-specific means were obtained by shifting the baseline by the predefined log2FC for that gene. Expression measurements were sampled from a normal distribution with gene-specific variance. To mimic missingness in scProteomics, each simulated measurement was masked as missing with a gene-specific probability based on our experimental data. Finally, the simulated dataset was analyzed with the limma linear model after filtering genes with sufficient observations (≥3 detected values per group), enabling DE detection performance to be evaluated relative to the known ground-truth log2FCs.

For data pre-processing, single cell runs were excluded if the number of identified proteins was below 1.5 or above 3 s.d. from cell line identification median.

For UMAP analysis, features with three or more measured values in all 6 groups were included; features with 50% or more measured values in one cell line and all missing values in another cell line were included to incorporate cell-line-specific missingness; within cell line, features with three or more measured values in one condition and all missing values in the other were included to incorporate condition-aware missingness to make a matrix of 1703 features and 32 cells. The missing values were replaced with zeros and the values were z-score scaled followed by log2 transformation.

For differential expression analysis within cell line, features with three or more measured values in both conditions were included without imputation; features with at most one missing value in one condition and all missing values in the other were included with imputation of column-wise minimum in the condition with all missing values. limma (v3.62.2) were used for statistical testing, genes with absolute log2FC larger than 0.58 and raw p value less than 0.1 were considered differentially expressed.

For the Venn diagram, the up-regulated proteins from each cell line were plotted as vertices in a network and connected with edges if they formed a co-complex or binary interaction pair in the HINT database. Proteins connected across cell lines were counted as overlaps in the Venn diagram. The empirical p value was calculated by shuffling the up-regulated proteins in the pool of the identified proteins.

## Author Contributions

Y.W. and Q.Z. contributed equally to this work.

## REFERENCES

(1) Kune, C.; Tielens, S.; Baiwir, D.; Fléron, M.; Vandormael, D.; Eppe, G.; Nguyen, L.; Mazzucchelli, G. SIGNIFICANT IMPACT OF CONSUMABLE MATERIAL AND BUFFER COMPOSITION FOR LOW-CELL NUMBER PROTEOMIC SAMPLE PREPARATION. Anal. Chem. 2025, 97 (7), 3836–3845. 10.1021/acs.analchem.4c03709.

(2) Zhang, Z.; Gao, Y. Evaluation of the Binding Preference of Microtubes for Nanoproteomics Sample Preparation. J. Proteome Res. 2023, 22 (1), 279–284. 10.1021/acs.jproteome.2c00477.

(3) Wang, Y.; Woo, J.; Sun, Z.; Assis, D.; Kirsch, Z.; Willetts, M.; Albano, M.; Liu, H.; Pienta, K. J.; Amend, S. R.; Zhang, H. Liquid Chromatographic and Mass Spectrometric Methods for Quantitative Proteomic Analysis from Single-Cell and Nanogram-Level Samples. Anal. Chem. 2025. 10.1021/acs.analchem.5c02808.

(4) Wallmann, G.; Leduc, A.; Slavov, N. Data-Driven Optimization of DIA Mass Spectrometry by DO-MS. J. Proteome Res. 2023, 22 (10), 3149–3158. 10.1021/acs.jproteome.3c00177.

(5) Krull, K. K.; Ali, S. A.; Krijgsveld, J. Enhanced Feature Matching in Single-Cell Proteomics Characterizes IFN-γ Response and Co-Existence of Cell States. Nat Commun 2024, 15 (1), 8262. 10.1038/s41467-024-52605-x.

(6) Yu, F.; Teo, G. C.; Kong, A. T.; Fröhlich, K.; Li, G. X.; Demichev, V.; Nesvizhskii, A. I. Analysis of DIA Proteomics Data Using MSFragger-DIA and FragPipe Computational Platform. Nat Commun 2023, 14 (1), 4154. 10.1038/s41467-023-39869-5.

(7) Budnik, B. SCoPE-MS: Mass Spectrometry of Single Mammalian Cells Quantifies Proteome Heterogeneity during Cell Differentiation. Genome Biology 2018, 19, 161–173.

(8) Fulcher, J. M.; Markillie, L. M.; Mitchell, H. D.; Williams, S. M.; Engbrecht, K. M.; Degnan, D. J.; Bramer, L. M.; Moore, R. J.; Chrisler, W. B.; Cantlon-Bruce, J.; Bagnoli, J. W.; Qian, W.-J.; Seth, A.; Paša-Tolić, L.; Zhu, Y. Parallel Measurement of Transcriptomes and Proteomes from Same Single Cells Using Nanodroplet Splitting. Nat Commun 2024, 15 (1), 10614. 10.1038/s41467-024-54099-z.

(9) Specht, H.; Emmott, E.; Petelski, A. A.; Huffman, R. G.; Perlman, D. H.; Serra, M.; Kharchenko, P.; Koller, A.; Slavov, N. Single-Cell Proteomic and Transcriptomic Analysis of Macrophage Heterogeneity Using SCoPE2. Genome Biol 2021, 22 (1), 50–77. 10.1186/s13059-021-02267-5.

(10) Liu, Y.; DiStasio, M.; Su, G.; Asashima, H.; Enninful, A.; Qin, X.; Deng, Y.; Nam, J.; Gao, F.; Bordignon, P.; Cassano, M.; Tomayko, M.; Xu, M.; Halene, S.; Craft, J. E.; Hafler, D.; Fan, R. High-Plex Protein and Whole Transcriptome Co-Mapping at Cellular Resolution with Spatial CITE-Seq. Nat Biotechnol 2023, 41 (10), 1405–1409. 10.1038/s41587-023-01676-0.

(11) Brunner, A.-D. Ultra-High Sensitivity Mass Spectrometry Quantifies Single-Cell Proteome Changes upon Perturbation. Molecular Systems Biology 2022, 18, 26–41.

(12) Ctortecka, C.; Clark, N. M.; Boyle, B. W.; Seth, A.; Mani, D. R.; Udeshi, N. D.; Carr, S. A. Automated Single-Cell Proteomics Providing Sufficient Proteome Depth to Study Complex Biology beyond Cell Type Classifications. Nat Commun 2024, 15 (1), 5707. 10.1038/s41467-024-49651-w.

(13) Reichmann, D.; Voth, W.; Jakob, U. Maintaining a Healthy Proteome during Oxidative Stress. Molecular Cell 2018, 69 (2), 203–213. 10.1016/j.molcel.2017.12.021.

(14) Leduc, A. Exploring Functional Protein Covariation across Single Cells Using nPOP. Genome Biology 2022, 23, 261–294.

(15) Woo, J.; Williams, S. M.; Markillie, L. M.; Feng, S.; Tsai, C.-F.; Aguilera-Vazquez, V.; Sontag, R. L.; Moore, R. J.; Hu, D.; Mehta, H. S.; Cantlon-Bruce, J.; Liu, T.; Adkins, J. N.; Smith, R. D.; Clair, G. C.; Pasa-Tolic, L.; Zhu, Y. High-Throughput and High-Efficiency Sample Preparation for Single-Cell Proteomics Using a Nested Nanowell Chip. Nat Commun 2021, 12 (1), 6246. 10.1038/s41467-021-26514-2.

(16) Zhu, Y.; Piehowski, P. D.; Zhao, R.; Chen, J.; Shen, Y.; Moore, R. J.; Shukla, A. K.; Petyuk, V. A.; Campbell-Thompson, M.; Mathews, C. E.; Smith, R. D.; Qian, W.-J.; Kelly, R. T. Nanodroplet Processing Platform for Deep and Quantitative Proteome Profiling of 10–100 Mammalian Cells. Nat Commun 2018, 9 (1), 882. 10.1038/s41467-018-03367-w.

(17) Derks, J. Increasing the Throughput of Sensitive Proteomics by plexDIA. Nature Biotechnology 2023, 41, 50–59.

(18) Liang, Y.; Truong, T.; Saxton, A. J.; Boekweg, H.; Payne, S. H.; Van Ry, P. M.; Kelly, R. T. HyperSCP: Combining Isotopic and Isobaric Labeling for Higher Throughput Single-Cell Proteomics. Anal. Chem. 2023, 95 (20), 8020–8027. 10.1021/acs.analchem.3c00906.

(19) Petrosius, V.; Aragon-Fernandez, P.; Arrey, T. N.; Woessman, J.; Üresin, N.; De Boer, B.; Su, J.; Furtwängler, B.; Stewart, H.; Denisov, E.; Petzoldt, J.; Peterson, A. C.; Hock, C.; Damoc, E.; Makarov, A.; Zabrouskov, V.; Porse, B. T.; Schoof, E. M. Quantitative Label-Free Single-Cell Proteomics on the Orbitrap Astral MS. July 31, 2024. 10.1101/2024.07.31.605978.

(20) 10x Genomics. IBT-16plex-Based Quantitative Proteomics at the Single-Cell Level: Enabling Protein Profiles toward the Single Cells from Mouse Spleen. 10k Human K562-r Cells, Singleplex Sample (Next GEM) — Count matrix filtered feature-barcode (H5) [Data set]. 10x Genomics. https://www.10xgenomics.com/datasets/10k-human-k562-r-cells-singleplex-sample-1-standard 2022.

(21) Mund, A. Unbiased Spatial Proteomics with Single-Cell Resolution in Tissues. Molecular Cell 2022, 82, 2335–2349.

(22) Vieth, B.; Ziegenhain, C.; Parekh, S.; Enard, W.; Hellmann, I. powsimR: Power Analysis for Bulk and Single Cell RNA-Seq Experiments. Bioinformatics 2017, 33 (21), 3486–3488. 10.1093/bioinformatics/btx435.

(23) Lai, W. K. M.; Mariani, L.; Rothschild, G.; Smith, E. R.; Venters, B. J.; Blanda, T. R.; Kuntala, P. K.; Bocklund, K.; Mairose, J.; Dweikat, S. N.; Mistretta, K.; Rossi, M. J.; James, D.; Anderson, J. T.; Phanor, S. K.; Zhang, W.; Zhao, Z.; Shah, A. P.; Novitzky, K.; McAnarney, E.; Keogh, M.-C.; Shilatifard, A.; Basu, U.; Bulyk, M. L.; Pugh, B. F. A ChIP-Exo Screen of 887 Protein Capture Reagents Program Transcription Factor Antibodies in Human Cells. Genome Res. 2021, 31 (9), 1663–1679. 10.1101/gr.275472.121.

